# Domestication reduces drought tolerance in watermelon through loss of root plasticity traits

**DOI:** 10.64898/2026.02.11.705227

**Authors:** Or E. shemer, Zvi M. bloom, Shira Gal, Gadi Peleg, Amnon Cochavi

## Abstract

Wild plants, particularly those native to xeric environments, are highly adapted to survive under harsh conditions. These adaptive strategies primarily ensure the successful transfer of genetic material to subsequent generations, often independently of fruit size or quality. In contrast, more than 10,000 years of domestication have shifted plant strategies away from survival-oriented traits toward increase in yield and fruit quality. In this study, we characterized both shared and divergent physiological traits contributing to drought tolerance in wild and domesticated watermelon genotypes. Specifically, we compared above- and belowground responses to water limitation in desert watermelon (*Citrullus colocynthis*) versus these in a watermelon cultivar (*Citrullus lanatus*). While aboveground responses to water scarcity were largely similar between the two genotypes, pronounced differences emerged belowground. Root biomass and surface area in the cultivated watermelon were predominantly concentrated in the upper soil layers. In contrast, desert watermelon displayed substantial root system plasticity under drought conditions. Although total root biomass remained largely distributed in the upper soil layers, root surface area shifted toward deeper soil layers, indicating enhanced water acquisition from deeper soil layers without additional biomass investment. These findings suggest that domesticated watermelon, despite originating from desert-adapted ancestors, has largely lost the capacity for dynamic root system adjustment in response to spatial and temporal variation in soil water availability.

## 1. Introduction

Over the past 10,000 years, plant domestication has largely favored traits that maximize reproductive output under managed agricultural systems. As a consequence, traits essential for the survival od domesticated plants in harsh environmental conditions, including drought and other climatic extremes, have been diminished (Bacher et al., 2022; Lupo & Moshelion, 2024; Wild et al. 2024). Sweet watermelon (*Citrullus lanatus*) originated in the arid regions of West Africa, where it has been domesticated for over 4,000 years (Chomicki & Renner, 2015; Paris, 2015). It was subsequently cultivated in the Mediterranean basin for approximately 2,000 years before spreading globally to Europe and China (Paris, 2015). In contrast, desert watermelon (*C. colocynthis*) originated in the arid regions of North Africa and remains widely distributed across sandy desert soils of the Middle East and central and southwestern Asia (Chomicki et al., 2020). Although *C. colocynthis* has long been used as a medicinal plant, it is not known to be fully domesticated (Levi et al., 2017; Chomicki et al., 2020). Notably, *C. colocynthis* plants collected in the wild exhibit substantial drought tolerance and are a considered to be a valuable genetic resource for enhancing drought tolerance in watermelon cultivars (*C. lanatus*), as the two species share the same chromosome number (2*n* = 22), have similar genome sizes, and are readily crossed with each other (Levi et al., 2017).

Plants naturally adapted to xerophytic conditions (including short rainy seasons, low soil water retention, and high vapor pressure deficit; VPD) possess a range of innate mechanisms that enable them to withstand drought stress. These mechanisms include: 1) Regulation of diffusive resistance to limit water loss. This strategy is primarily mediated by stomatal regulation to reduce water loss through transpiration under water-limiting conditions (Gillon & Yakir, 2000). Plant species exhibit stomatal behavior ranging from early stomatal closure during water deficit (isohydric) to stomata remaining open despite declining in water availability (anisohydric). However, according to Hochberg et al. (2018), no species is strictly isohydric or anisohydric and the threshold for stomatal closure varies among species and stress intensities. While stomatal closure limits internal CO₂ concentration, some plant species are able to sustain sufficient levels of carbon assimilation by increasing intrinsic water-use efficiency (Jones, 1998; Zait & Schwartz, 2018; Cochavi et al., 2021). 2) Modification of shoot architecture to reduce transpiration. Here, many xerophytic plants develop small succulent leaves to reduce transpirational surface area. A low shoot-to-root ratio further contributes to maintaining water balance under extreme drought conditions (Bloom et al., 1985). Additionally, increased leaf and cuticle thickness limits non-stomatal water loss and enhances drought resistance (Aneja et al., 2025). 3) Development of a robust root system to enhance water acquisition. Xerophytic plants often develop extensive root systems to maximize water uptake from different soil layers. Drought stress reduces root’s water and nutrient uptake and translocation to aerial tissues. Expansion of root biomass, regulated by specific genetic pathways (Uga et al., 2013; Marques et al., 2025) enhances water absorption under water-deficient conditions (Zhao et al., 2017). Accordingly, overall root system size (root diameter, root length, and fibrous root capacity) is positively associated with plant performance under drought stress (Arsova et al., 2019). The elongation capacity of root system is a critical trait under unstable rainfall regimes. This is since it facilitates access to water and nutriant sources across different soil layers (Lozano et al., 2020). Plant species develop distinct strategies adapted to their endemic environment, while csrtain plant species develop deep root systems to access groundwater (Uga et al., 2013; Kwatcho Kengdo et al., 2025)others produce extensive lateral roots in shallow layers to absorb runoff water (Nakhforoosh et al., 2021).

The composition of the root system plays a critical role in plant resilience to drought. Fine roots are particularly important for water and nutrient uptake (McCormack et al., 2015; Cochavi et al., 2020). The high specific root length (SRL; total root length per unit biomass, m g⁻¹) and specific root surface area (SRA; total root surface area per unit biomass, m² g⁻¹) of fine roots facilitate efficient water absorption from the surrounding rhizosphere (Comas et al., 2013). In addition to their low carbon cost and short lifespan, fine roots allow the plant to adjust to water limitation without substantially diverting energy from aboveground growth (Hou et al., 2024). Moreover, their small diameter, together with the exudation of sugars and mucilage, enables fine roots to maintain close contact with soil particles even as the soil shrinks during drought, thereby sustaining water flow from the soil to the plant (Carminati & Vetterlein, 2013; Xiao et al., 2020). Overall, root trait plasticity plays a pivotal role in plant adaptation to shifts in environmental conditions (Zhu et al., 2010; Barberon et al., 2016; Zhao et al., 2017). Excessive root branching in well-watered zones or in soil layers that contribute little to water uptake can result in inefficient allocation of carbon from the shoot to the root system. Precise spatial and temporal regulation of root traits is essential for optimizing plant performance under drought stress.

Recent advances in research and technology have enabled accurate characterization and functional interpretation of plant root systems (Tracy et al., 2019; Zhang et al., 2025). Numerous genes associated with root architecture and dynamics have been identified, elucidating the roles of root systems in drought tolerance (Bengough et al., 2011; Uga et al., 2013; Marques et al., 2025). In parallel, root phenotyping techniques have advanced substantially, and modern tools now enable accurate three-dimensional characterization of root system architecture (Smith et al., 2022; Govta et al., 2025; Michels et al., 2025).

In the current study, we aim to characterize physiological and morphological differences between domesticated and wild watermelon under drought conditions. By integrating multiple analytical approaches, we seek to quantify whole-plant responses to dynamic changes in soil water availability.

## Material and methods

### 2.1 Plant material and growing conditions

Watermelon seeds obtained from the Agricultural Research Organization (ARO) Germplasm Collection at the Newe Ya’ar Research Center (Ramat Yishai, Israel) were used in this study. The cultivar ‘Sugar Baby’ (*C. lanatus*) served as the domesticated watermelon, while a wild *C. colocynthis* accession collected in Hazeva (30°46′28″N, 35°14′20″E), located in the Arava Valley, southern Israel, was used as the desert watermelon and numbered 16207. Plants were grown in two soil types-fine-clayey and sandy-and in two container systems: 4-L pots and cylindrical tubes (1 m depth × 20 cm diameter). The 4-L pots were filled with a high water-retention clayey soil (chromic Haploxerert, fine-clayey, montmorillonitic, thermic), consisting of 55% clay, 25% silt, 20% sand, and 2% organic matter (pH 7.2), to minimize rapid soil dehydration. In contrast, the cylindrical tubes were filled with pure sandy soil to prevent waterlogging within the root zone. All experiments were conducted in a net house under ambient environmental conditions during the spring season (April–June).

### 2.2 Pots experiment

To evaluate gas exchange responses to dehydration, seeds of the two accessions, ‘Sugar Baby’ (*C. lanatus*) and the Hazeva line (*C. colocynthis*), were sown separately in five 4-L pots per accession and arranged in a randomized design within the net house. The dehydration experiment was initiated after establishment of young plants at the 4-5 true-leaf stage. Gas exchange measurements were conducted every 2-3 days using a LI-6800F portable gas exchange and chlorophyll fluorescence system (LI-COR Biosciences, Lincoln, NE, USA). Measurement conditions within the LI-6800F chamber were controlled as follows: CO₂ concentration of 410 ppm, photosynthetic photon flux density of 1,000 µmol m⁻² s⁻¹, relative humidity of 40%, and chamber temperature set to 25 °C, approximating ambient conditions. During each measurement session, relative soil water content in each pot was assessed using a true time-domain reflectometer (TDR; Acclima Inc.), following a two-point calibration (0 and 100%) performed weekly. Leaf and root samples were collected for anatomical analyses at the conclusion of the experiment (see Plant histology section).

To evaluate the dynamics of root system responses to drought, a continuous drought experiment was conducted under controlled conditions. As in the previous experiment, ‘Sugar Baby’ and the Hazeva line seeds were sown in 4-L pots filled with clayey soil. After plant establishment at the 4-5 true-leaf stage, half of the plants in each group were subjected to dehydration treatment. Once per week, four plants from each genotype × treatment combination were measured using a LI-600 porometer/fluorometer (LI-COR Biosciences) over a four-week period, resulting in a total of 16 pots per genotype × treatment. Following the final measurement, shoots were excised and oven-dried at 72 °C for 48 h to determine shoot dry weight. The soil was then gently washed from the root systems, which were subsequently scanned at 800 dpi using the WinRHIZO® system (Regent Instruments Inc., Ottawa, ON, Canada). Roots were then oven-dried at 72 °C for 48 h, and root dry weight was recorded for calculation of root-to-shoot (R/S) ratios.

### 2.3 Cylindrical Tube experiment

Plants were grown in cylindrical plastic tubes (100 cm depth × 20 cm diameter) filled with a clean sand medium. A high-density mesh screen was installed at the bottom of each tube to allow drainage and minimize the risk of waterlogging during root architecture assessment. Small holes were drilled along the tube walls at 25 cm intervals, starting 12.5 cm above the bottom, to enable soil water content measurements using time-domain reflectometry (TDR). Seeds of ‘Sugar Baby’ and the Hazeva line were sown in the tubes, with six biological replicates per genotype. After plant establishment at the four–five true-leaf stage, irrigation was withheld from half of the plants in each group. Once per week, porometer measurements were conducted on the youngest fully expanded leaf. Soil water content was subsequently measured at four depths along the soil profile. The experiment was terminated once photosynthetic efficiency (Y(II)) stabilized at minimal values, prior to the onset of leaf senescence. Shoots were harvested, and leaves were oven-dried at 72 °C for 48 h to determine dry biomass. The sand was then carefully removed from the tubes and divided into depth segments (0–25, 25–50, 50–75, and 75–100 cm below the soil surface). Roots from each depth segment were gently washed free of sand, scanned using the WinRHIZO® system, and subsequently oven-dried at 72 °C for 48 h to determine root dry weight.

### 2.4 Root phenotypic evaluation

Root system spatial distribution under drought was evaluated using 10-L pots filled with clean sand. Seeds of ‘Sugar Baby’ and the Hazeva line were sown with ten biological replicates per accession. Due to non-uniform seedling emergence, nine ‘Sugar Baby’ plants and seven Hazeva plants were finally included in the experiment. After establishment of young plants at the 4-5 true-leaf stage, the drought treatment was initiated using four plants of ‘Sugar Baby’ and four plants of the Hazeva line. Plants were grown until drought-treated individuals exhibited daytime loss of leaf turgor. At this point, root phenotyping was performed using the Phenoroot scanning system (Rehovot, Israel; https://www.phenoroot.com), which generated 24 root architectural parameters. These parameters were subsequently analyzed using the RhizoVision Explorer software (Seethepalli et al., 2020).

### 2.5 plant histology

For anatomical analyses, the youngest fully expanded leaves of ‘Sugar Baby’ and the Hazeva line were excised into ∼5 mm² segments and fixed for 48 h in FAA solution (10% formaldehyde, 5% acetic acid, and 50% ethanol, v/v in water) at 4 °C in the dark. Fixation was followed by dehydration through a graded ethanol series (50%, 70%, 90%, 95%, 100%, and 100% × 2) and subsequent replacement of ethanol with Histoclear (xylene substitute). Samples were then embedded in paraffin, and 10-µm transverse sections were prepared using a Leica RM2245 microtome (Leica Biosystems, Nussloch, Germany). Sections were stained with safranin and fast green (Ruzin, 1999) and imaged under a light microscope (Leica DM500, Heerbrugg, Switzerland) equipped with an ICC50 HD camera. Leaf thickness, palisade and spongy mesophyll thickness, and cuticle thickness were quantified using ImageJ software.

Root anatomical sections were obtained from the main root (MR) and fine roots (FR). Main root sections were collected from the youngest region of the root, immediately proximal to the first branching point. Root fixation and staining followed the protocol described by Freschet et al. (2021), using a fixative composed of 2% glutaraldehyde (v/v, pH 6.8) and 2% formaldehyde (v/v, pH 6.8) in 0.1 M cacodylate buffer. Sections were subsequently stained with safranin and alcian blue (Wolberg et al., 2023). Quantitative anatomical traits, including root diameter, stele area, cortex area, xylem area, and mean xylem vessel diameter, were measured using ImageJ software.

### 2.6 Statistical analysis

To evaluate the intrinsic and water use efficiency (*A_n_* to *g_sw_* or E ratio, respectively), a curve fitted to the gas exchange measured data:

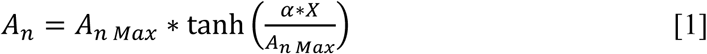

Where *A_nMax_* is the maximal calculated *A_n_* value, α is the slope of the linear phase of the curve, and X is the dependent variable (g_sw_, or E). The saturation point of the curve can be calculated:

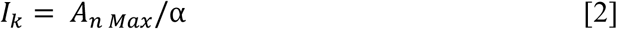

Equations 1&2 were also used to describe the SWC relation with the root system SA. The effect of the soil water content on the ratio between *A_n_* to the Electron Transport Rate (*ETR*) was described by exponential decay equation:

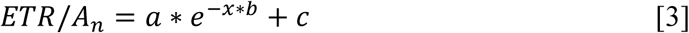

Where *a* is the maximal *ETR/A_n_* value, *b* is the decay coefficient, and *c* is the horizontal asymptote value.

All the equations coefficients were optimized through minimizing the Root Mean square Error (RMSE) of their variance using Excel Solver function:

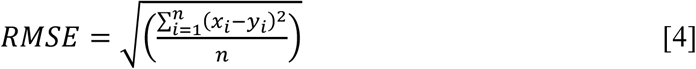

Where 𝑥_𝑖_ and 𝑦_𝑖_ are the measured and estimated values, respectively.

Statistical analyses were conducted using two-way ANOVA, followed by mean separation with Tukey’s HSD test (α < 0.05). A Bonferroni correction was applied to account for unequal sample sizes. Data normality was assessed using the Shapiro-Wilk test, and homogeneity of variances was verified using Levene’s test. When assumptions were violated, data were transformed prior to analysis, as specified in the Results section.

## 3. Results

### 3.1 Aboveground physiology is similar between genotypes

Comparison of water use efficiency (WUE) and intrinsic water use efficiency (iWUE) between the wild “Hazeva Line” (*C. colocynthis*) and domesticated ‘Sugar Baby’ watermelon (*C. lanatus*) showed similar relationships under drought conditions. The slope of the WUE response was comparable between genotypes, with values of 3.98 for the domesticated watermelon and 3.16 for the wild watermelon (Figure 1a; Table 1). A similar pattern was observed for iWUE, with slopes of 103.09 and 87.87 for the domesticated and wild watermelons, respectively (Figure 1b; Table 1). Accordingly, the saturation constants (*I_k_*) for both WUE and iWUE did not differ substantially between species. In contrast, the maximum net assimilation rate (An_max_), estimated from the WUE curve, was significantly higher in the domesticated ‘SB’ watermelon (30.84 µmol m⁻² s⁻¹) than in the wild watermelon (18.71 µmol m⁻² s⁻¹).

**Figure 1.**
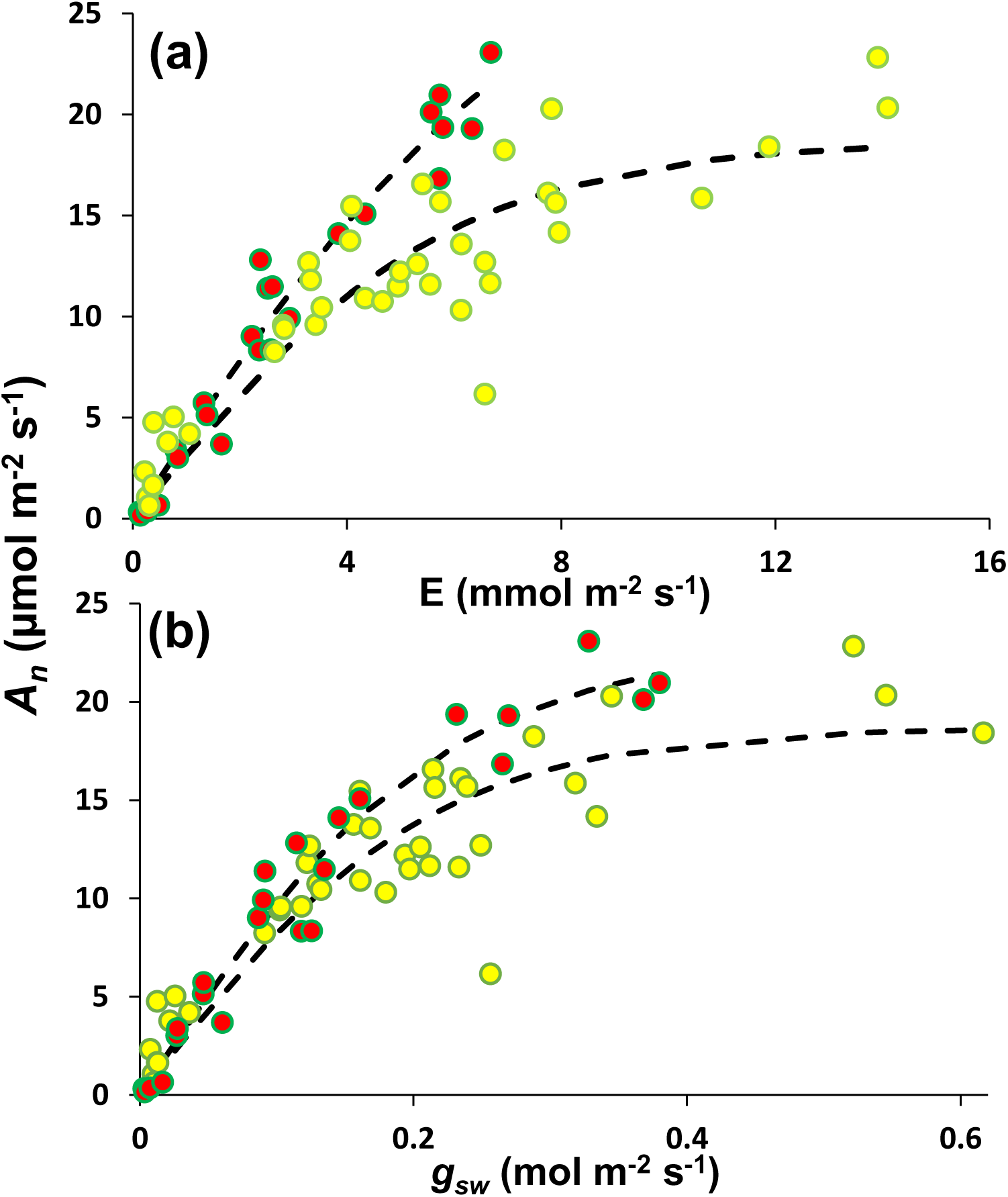
Relation between transpiration rate (*E*) or stomatal conductance (gsw) to carbon assimilation rate (*An*) in *C. lanatus* (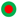) and *C. colocynthis* (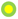). Fitted curves coefficients are attached in table 1.

**Table 1.** Coefficients of the fitted curves 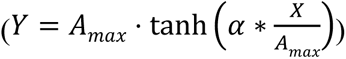 describing the relationships between carbon assimilation rate (An) and transpiration rate or stomatal conductance in figure XX. Curves were fitted by minimizing the RMSE. *Iₖ* denotes the X-axis value at which An reaches saturation.

| Specie | $A_n$-E | | | | $A_n$-$g_{sw}$ | | | |
| --- | --- | --- | --- | --- | --- | --- | --- | --- |
| | $A_{max}$ | $\alpha$ | $I_k$ | RMSE | $A_{max}$ | $\alpha$ | $I_k$ | RMSE |
| *C. lanatus* | 30.84 | 3.98 | 7.74 | 1.42 | 22.82 | 103.09 | 0.22 | 1.53 |
| *C. colocythis* | 18.71 | 3.16 | 5.92 | 2.56 | 18.68 | 87.87 | 0.21 | 2.79 |

The *ETR/An* ratio differed significantly between the accessions across varying soil water content (SWC) levels (Figure 2). Under well-watered conditions, the ratio (parameter c) was approximately 50% higher in the wild accession than in domesticated watermelon (15.31 vs. 10.45, respectively; Table 2). As soil moisture declined, *ETR/An* values rose sharply in both accessions. The calculated maximal *ETR/An* ratios reached 148.44 and 345.03 for the wild and domesticated watermelon, respectively, indicating a more pronounced decoupling of electron transport from carbon assimilation to respiration in the domesticated versus with the wild watermelon under severe water stress.

**Figure 2.**
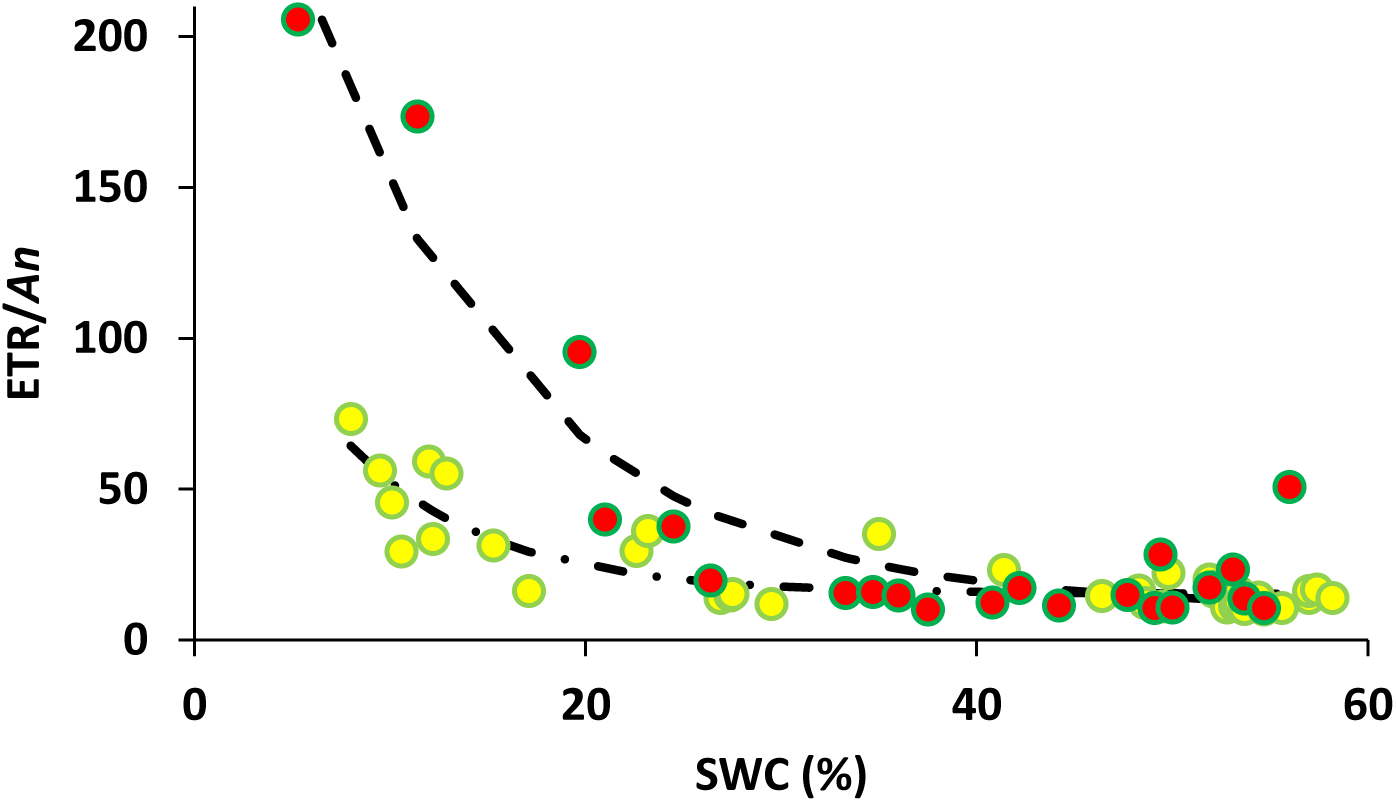
Relation between the measured volumetric soil water content (SWC) to the ratio between the electron transport rate and carbon assimilation (*ETR/An*) in *C. lanatus* (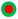) and *C. colocynthis* (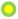). Fitted curves coefficients are attached in table 2.

**Table 2.**
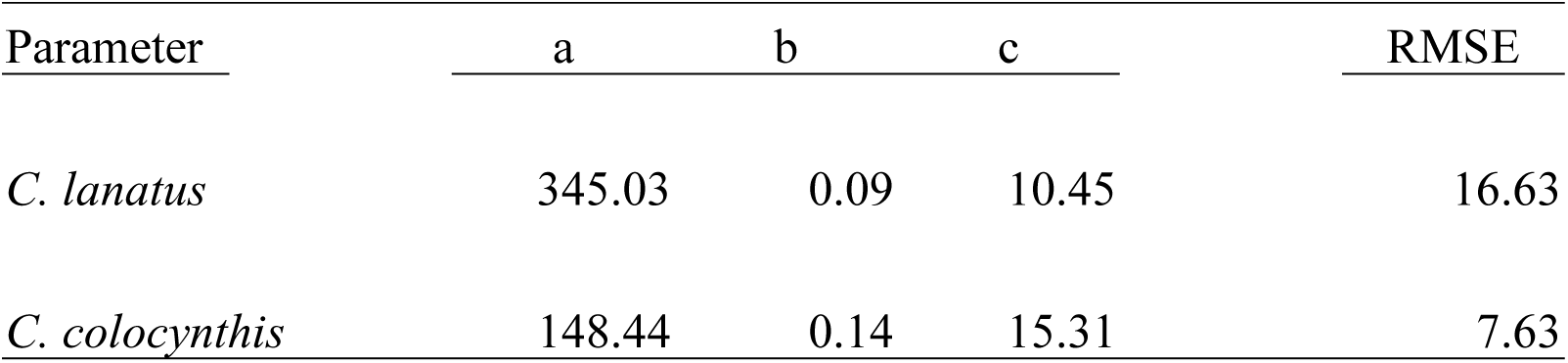
Coefficients of three parameters exponential decay equation y=𝑎∗𝑒^(−𝑥∗𝑏)+𝑐 describing the relation between the ETR/An ratio to the measured relative soil water content (SWC).

Histological analysis of leaf and root sections revealed a significant genotype effect, while irrigation treatment had no discernible influence on these parameters (Figure 3). Total leaf thickness was approximately 50% greater in the wild versus domesticated watermelon (∼150 vs. 100 µm; Figure 4a). Similarly, the cuticle thickness of the wild was double than that of the domesticated watermelon (16 vs. 8 µm; Figure 4b).

**Figure 3.** Leaf (upper panels) and main root (lower panels) cross sections of wild and domesticated watermelons.

**Figure 4.**
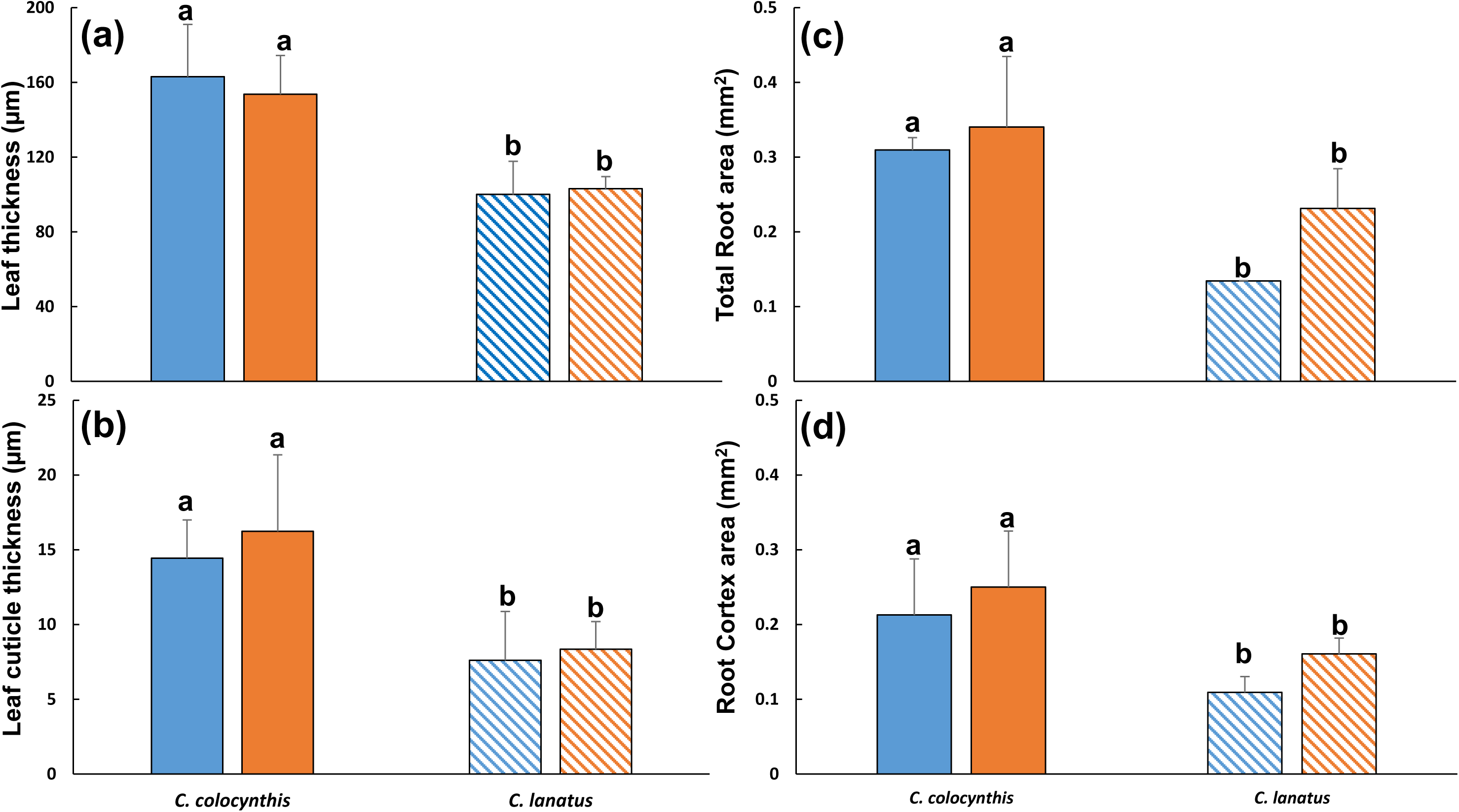
Histological analysis of wild and domesticated watermelon under control (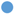) and drought (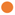) conditions. (a) leaf thickness and (b) cuticle thickness, measured in six replicates (*n*=6). (c) Total area of the main root sections, and (d) Cortex area were measured in three replicates (*n*=3). Different letters indicate statistically significant difference according to a Tukey-HSD test (α<0.05).

In contrast, fine root histology showed no significant differences across genotypes or irrigation treatments. However, the total cross-sectional area of the main root was approximately 50% larger in the wild watermelon (0.3 vs. 0.2 mm^2^; Figure 4c). A similar trend was observed for the cortex area (0.23 mm^2^ in the wild vs. 0.13 mm^2^ in the domesticated accession; Figure 4d). Notably, no significant differences were detected in the stele area between the two genotypes.

Below- and above ground growth dynamics were distinct between the domesticated and wild accessions (Figure 5). Although initial root surface areas were comparable, *C. colocynthis* (wild) developed a more extensive root system over the experimental period (Figure 5a). Specifically, the irrigated *C. colocynthis* was the only treatment to exhibit a continuous increase in root surface area throughout the study. By day 21, a significant divergence was observed between water treatments; root surface area in the domesticated accession was 649 cm^2^ (irrigated) vs. 198 cm^2^ (drought), while in the wild accession, it reached 1342 cm^2^ vs. 710 cm^2^, respectively (Table 3).

**Figure 5.**
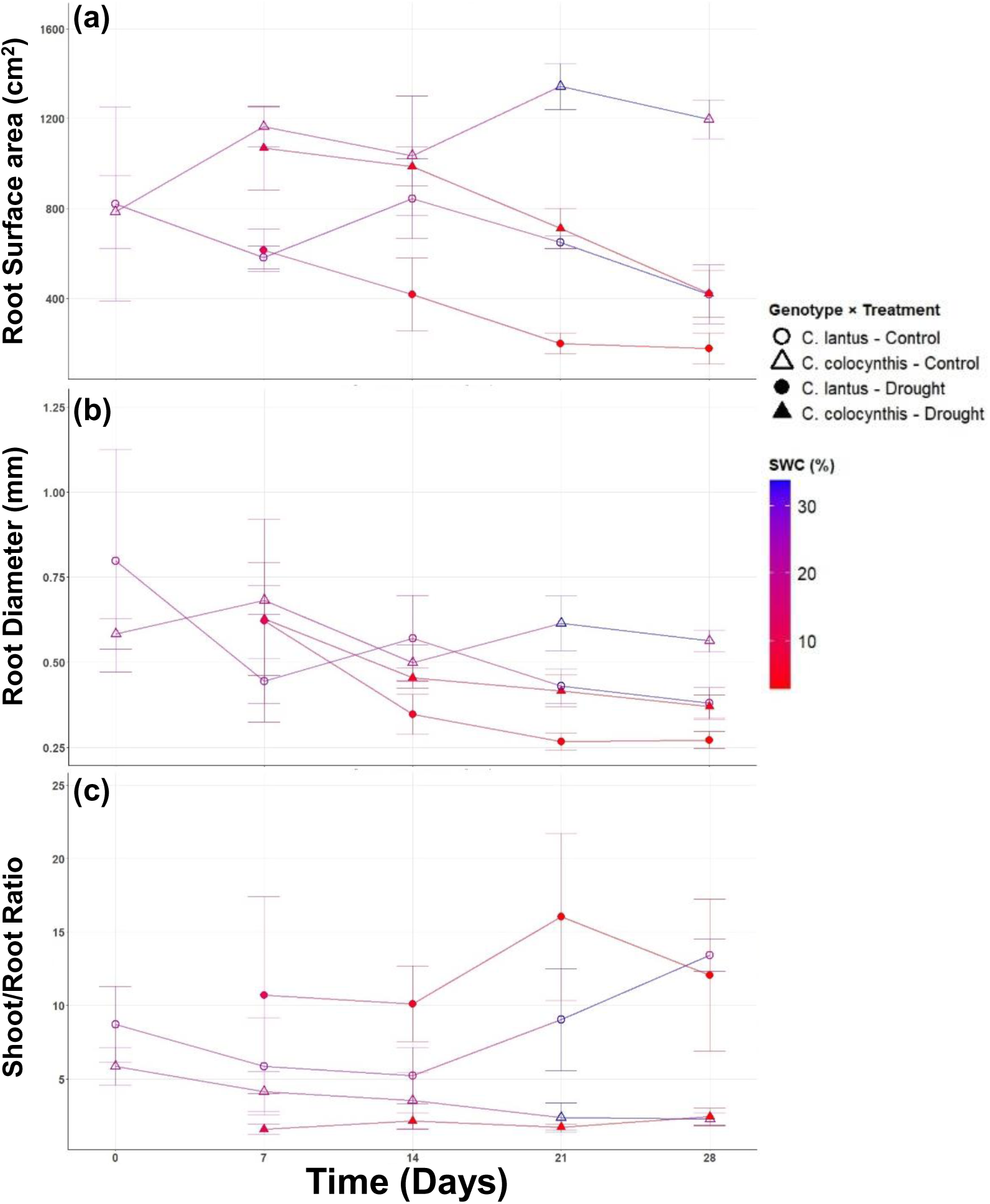
Dynamics of (a) root surface area, (b) root diameter, and (c) Shoot/Root ratio in wild and domesticated watermelons under control and drought conditions. The color gradient represents the mean soil water content (SWC) for each treatment. Each data point represents the mean of four replicates (*n*=4); error bars indicate ±SD. Detailed statistical analysis is provided in Table 3.

**Table 3.** Mean values (±SD) for the data presented in Figure 3. different letters within the same time point indicate statistically significant differences according to Tukey-HSD test (α<0.05; *n*=4).

| Trait | Time (Days) | *C.Colocynthis* |  | *C.lanatus* |  |
| --- | --- | --- | --- | --- | --- |
|  |  | Control | Drought | Control | Drought |
| Surface area (cm<sup>2</sup>) | 0 | 785.39±162.66a | | 820.81±431.39a | |
| | 7 | 1163.52±87.18a | 1068.73±185.90a | 582.35±51.19b | 614.72±95.01b |
| | 14 | 1035.02±265.10a | 986.35±86.84a | 843.04±177.24ab | 416.68±162.88b |
| | 21 | 1342.97±102.84a | 710.79±90.5b | 649.2±27.07b | 198.95±47.31c |
| | 28 | 1197.02±86.62a | 420.51±105.47b | 417.22±131.22b | 177.32±69.59c |
| Diameter (mm) | 0 | 0.58±0.04a | | 0.79±0.32a | |
| | 7 | 0.68±0.04a | 0.62±0.106a | 0.44±0.06a | 0.62±0.29a |
| | 14 | 0.49±0.05ab | 0.45±0.02ab | 0.57±0.12a | 0.34±0.05b |
| | 21 | 0.61±0.07a | 0.41±0.04b | 0.43±0.05b | 0.26±0.02c |
| | 28 | 0.56±0.03a | 0.37±0.03b | 0.37±0.04b | 0.27±0.02c |
| Shoot/Root Ratio | 0 | 0.19±0.03a | | 0.12±0.04a | |
| | 7 | 0.27±0.09ab | 0.36±0.16a | 0.27±0.21ab | 0.18±0.11b |
| | 14 | 0.48±0.18ba | 0.45±0.07a | 0.23±0.11b | 0.10±0.03c |
| | 21 | 0.66±0.15a | 0.50±0.14a | 0.13±0.04b | 0.07±0.02b |
| | 28 | 0.59±0.05a | 0.43±0.091a | 0.07±0.01b | 0.09±0.02b |

Average root diameter trends mirrored those observed for surface area (Figure 5b). Both genotypes showed a significant reduction in root diameter under drought stress. While the average root diameter of irrigated *C. colocynthis* and *C. lanatus* was identical at 28 days (0.37 mm), drought-stressed plants diverged; *C. lanatus* exhibited the thinnest roots (0.27 mm), whereas irrigated *C. colocynthis* produced the thickest (0.56 mm).

The R\S ratio remained primarily genotype-dependent and was less sensitive to water status (Figure 5c). Throughout the experiment, *C. colocynthis* maintained a higher, more balanced R\S ratio, whereas the biomass of *C. lanatus* was consistently skewed toward shoot development.

Root surface area (SA) in both species was significantly influenced by soil water content (SWC; Figure 6). The maximal surface area (SA_max_) achievable within the four-liter pot volume was approximately 60% higher in *C. colocynthis* compared to *C. lanatus* (1119 vs. 681 cm^2^, respectively; Table 4). However, the rate of SA reduction in response to declining SWC (parameter *α*) was more than twice as rapid in *C. colocynthis* than in *C. lanatus* (82 vs. 203, respectively). Furthermore, *C. colocynthis* reached its maximal SA at a lower SWC threshold (*I_k_*) compared to *C. lanatus* (5.49 vs. 8.31).

**Figure 6.**
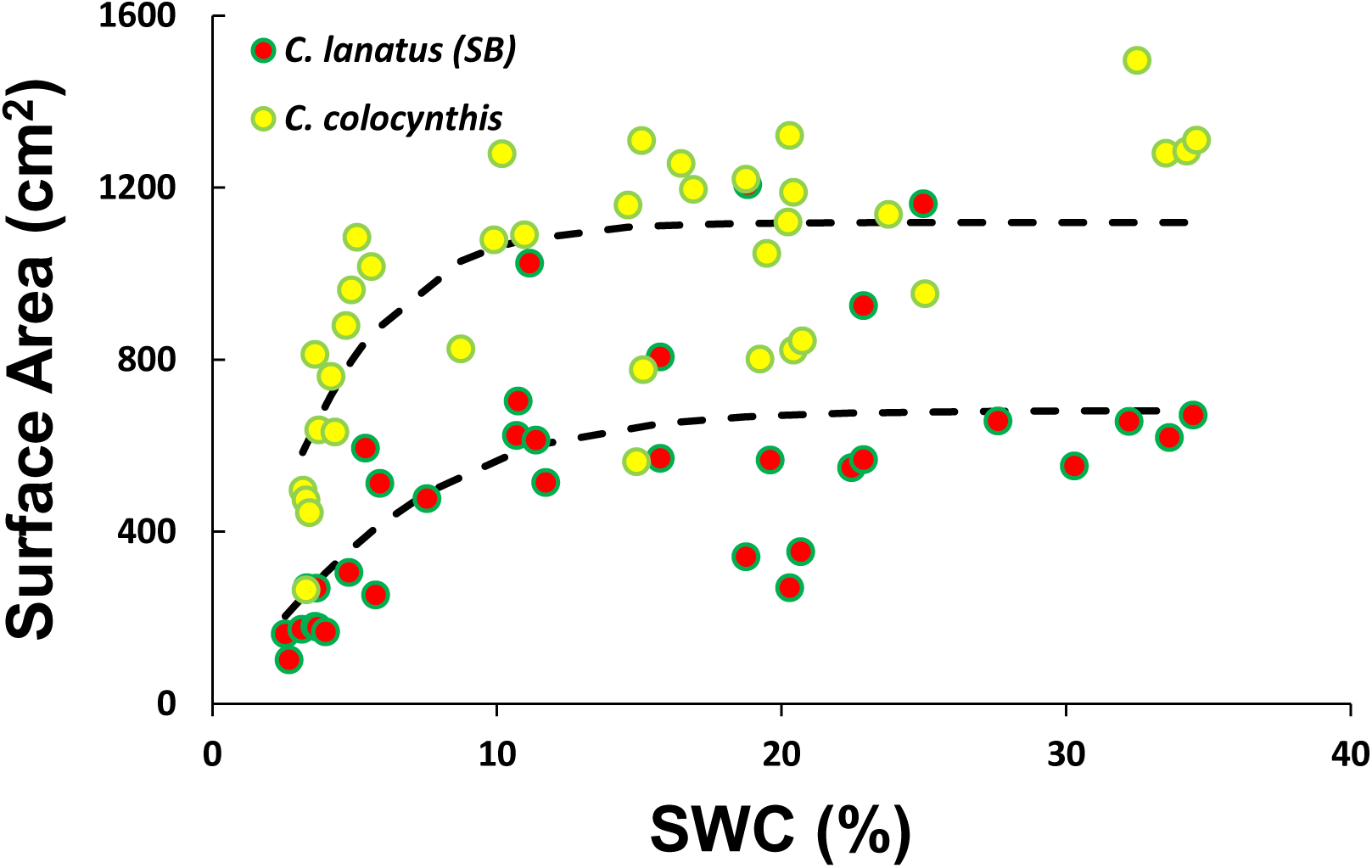
Relation between the measured volumetric soil water content (SWC) to the root surface area in *C. lanatus* (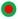) and *C. colocynthis* (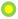). Fitted curves coefficients are attached in table 4.

**Table 4.**
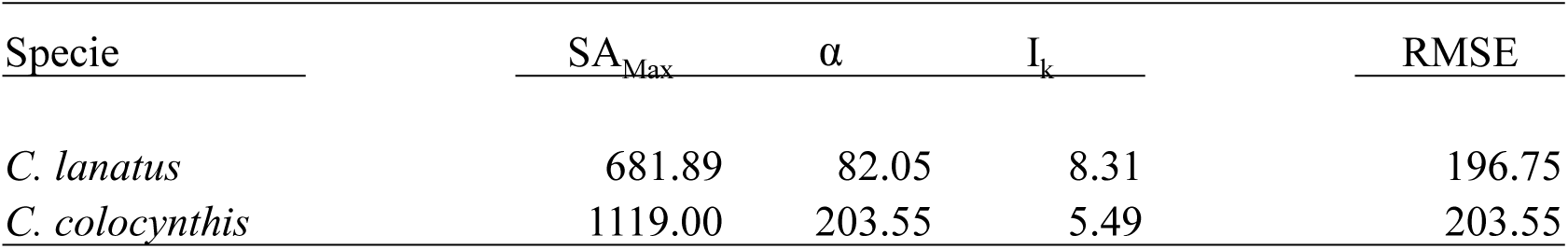
Coefficients of the fitted curves 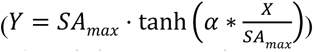 describing the relationships between Root Surface area (SA) and the measured volumetric soil water content (SWC) in figure XX. Curves were fitted by minimizing the RMSE. *Iₖ* denotes the X-axis value at which SA reaches saturation.

Watermelon genotypes were evaluated in a tube experiment to characterize the spatial distribution of their root systems under drought conditions. The SWC was monitored at four depths throughout the study to quantify stress levels (Figure S1). Over the course of the experiment, distinct photosynthetic responses to varying SWC were observed (Figure 7). In the wild accession, a negative relationship was found between Y(II) and increasing SWC under well-watered conditions (Figure 7a). Conversely, under drought conditions, Y(II) values remained high. The Y(II) response to increasing SWC was more moderate in the domesticated watermelon (slope of -0.01 compared to -0.07 in the wild accession); however, a significant reduction in Y(II) was observed under drought conditions (Figure 7b).

**Figure 7.**
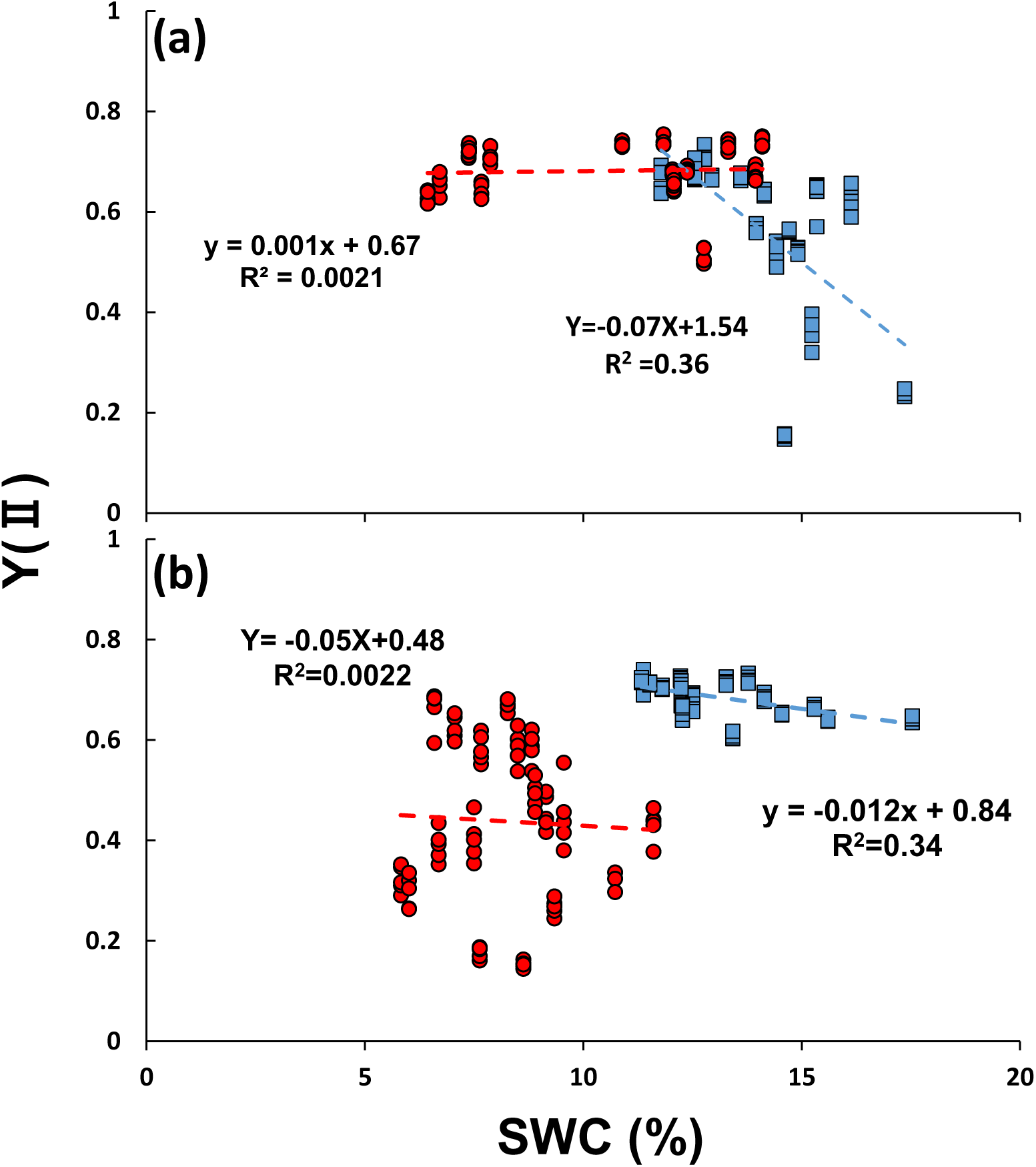
the effect of average soil water content on the photosynthetic efficiency (Y(II)) of (a) *C. colocynthis* and (b) *C. lanatus* under control (blue) and drought (red) conditions in one meter depth cylinder. Linear regression equations was added for each group (genotype*treatment).

The spatial distribution of the root system revealed divergent patterns in root SA between the wild and domesticated accessions, despite similar biomass distributions (Figure 8). In both species, the vertical distribution of root biomass remained relatively unchanged by the irrigation treatment (Figure 8a,b). The majority of the biomass, approximately 40% in the wild and 60% in the domesticated watermelon, was concentrated in the upper soil layer (0–25 cm), with a decreasing proportion in deeper layers. In contrast, the spatial distribution of root SA in response to drought differed significantly between the species (Figure 8c,d). In the wild watermelon, root SA distribution shifted under drought conditions; while the SA was primarily located in the upper soil layer under irrigation, it shifted significantly toward deeper soil layers under drought. Conversely, the SA distribution of the domesticated watermelon remained localized in the upper soil layer and was found to be inert to SWC status.

**Figure 8.**
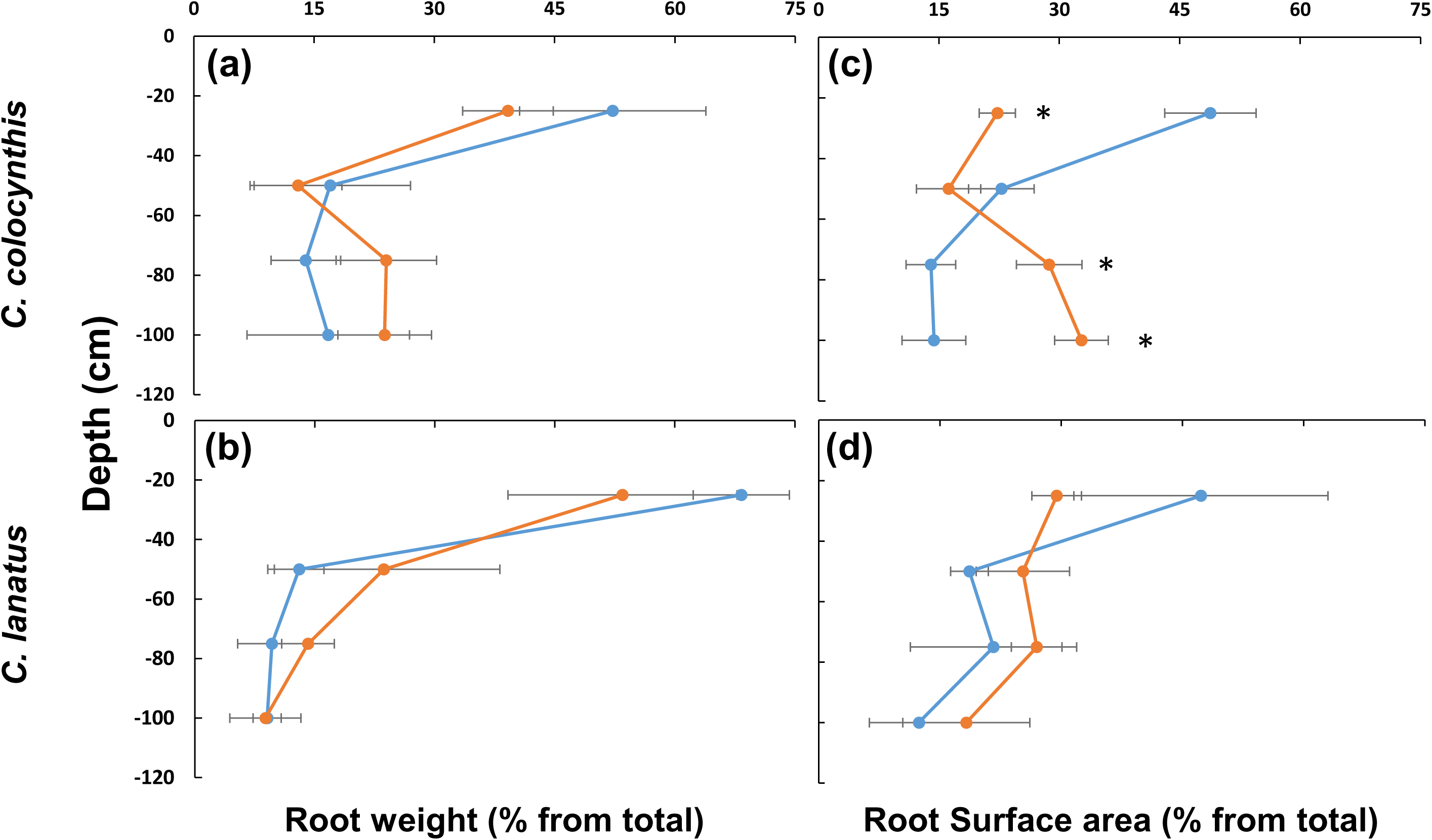
Effects of drought (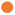), and irrigation (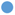) treatments on root biomass and surface area distribution of *C. colocynthis* and *C. lanatus* across soil depths. Left panels: relative root biomass, right panels: relative root surface area. Values represent mean ± SD (*n* = 3), asterisk represent statistical difference between treatments according to Tukey-HSD test(α<0.05).

Three-dimensional scanning of root systems was conducted for both species at the early growth stage, resulting in the quantification of 27 morphological parameters (Figure 9). Most of these parameters were significantly affected by water status (Figure S2). Root system depth was the only trait significantly influenced by both genotype and water status (Figure 10). Under well-watered conditions, both wild and domesticated watermelon accessions developed deeper root systems. Notably, the root depth of the wild watermelon under drought conditions was comparable to that of the irrigated domesticated variety. In addition, root system network area was significantly greater under irrigation in both genotypes (Figure 10b). However, under drought stress, a pronounced reduction in network area was observed only in the domesticated watermelon relative to its irrigated control. Analysis of root tip number revealed no significant differences between genotypes or irrigation treatments (Figure S3).

**Figure 9.**
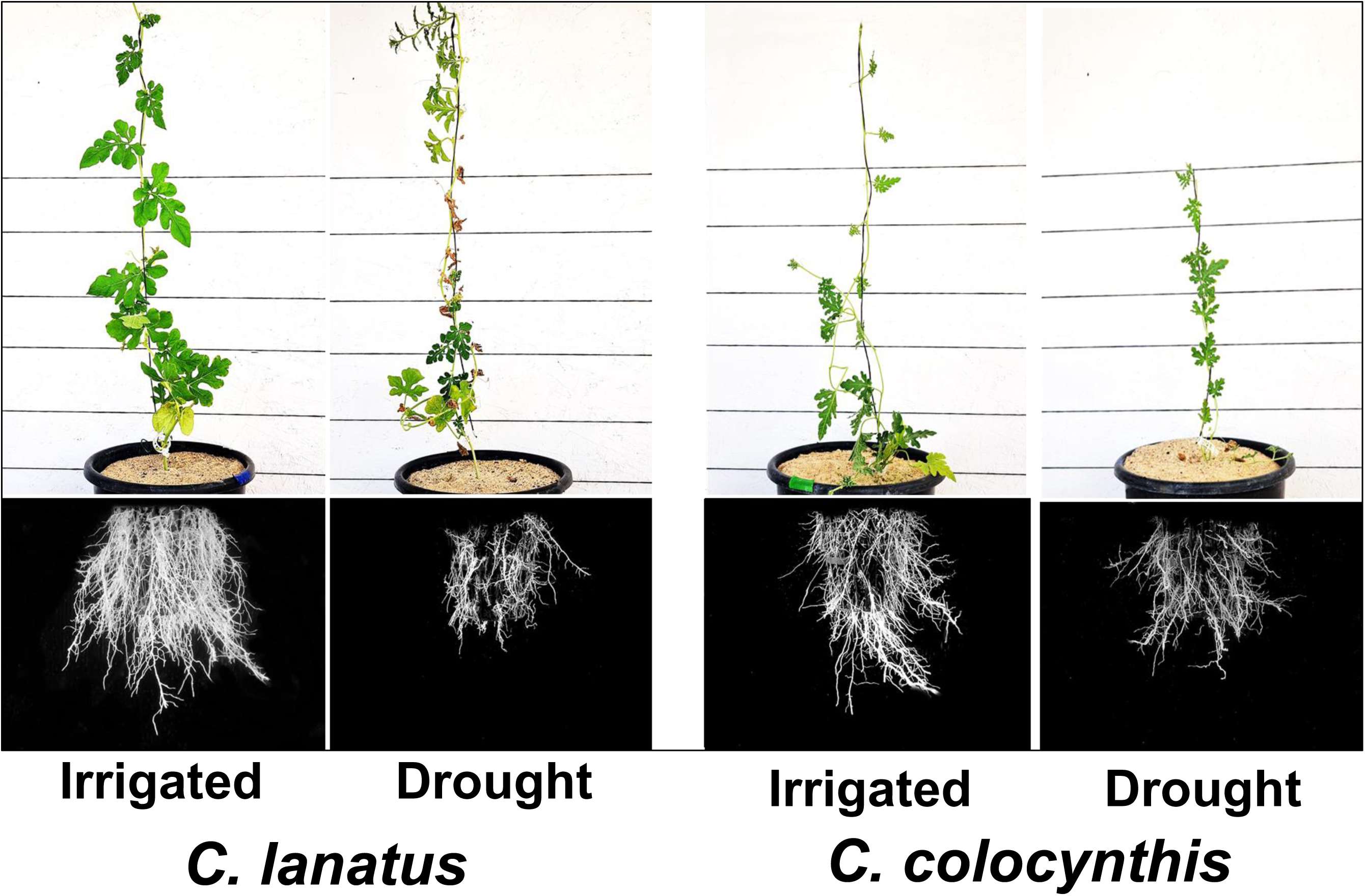
Demonstration of the outcome of the PhenoRoot® Root scan system. Plants were grown in 10 liter pots filled with clean sand and scanned after turgor loss in the drought treated plants. Statistical data attached in figure S2.

**Figure 10.**
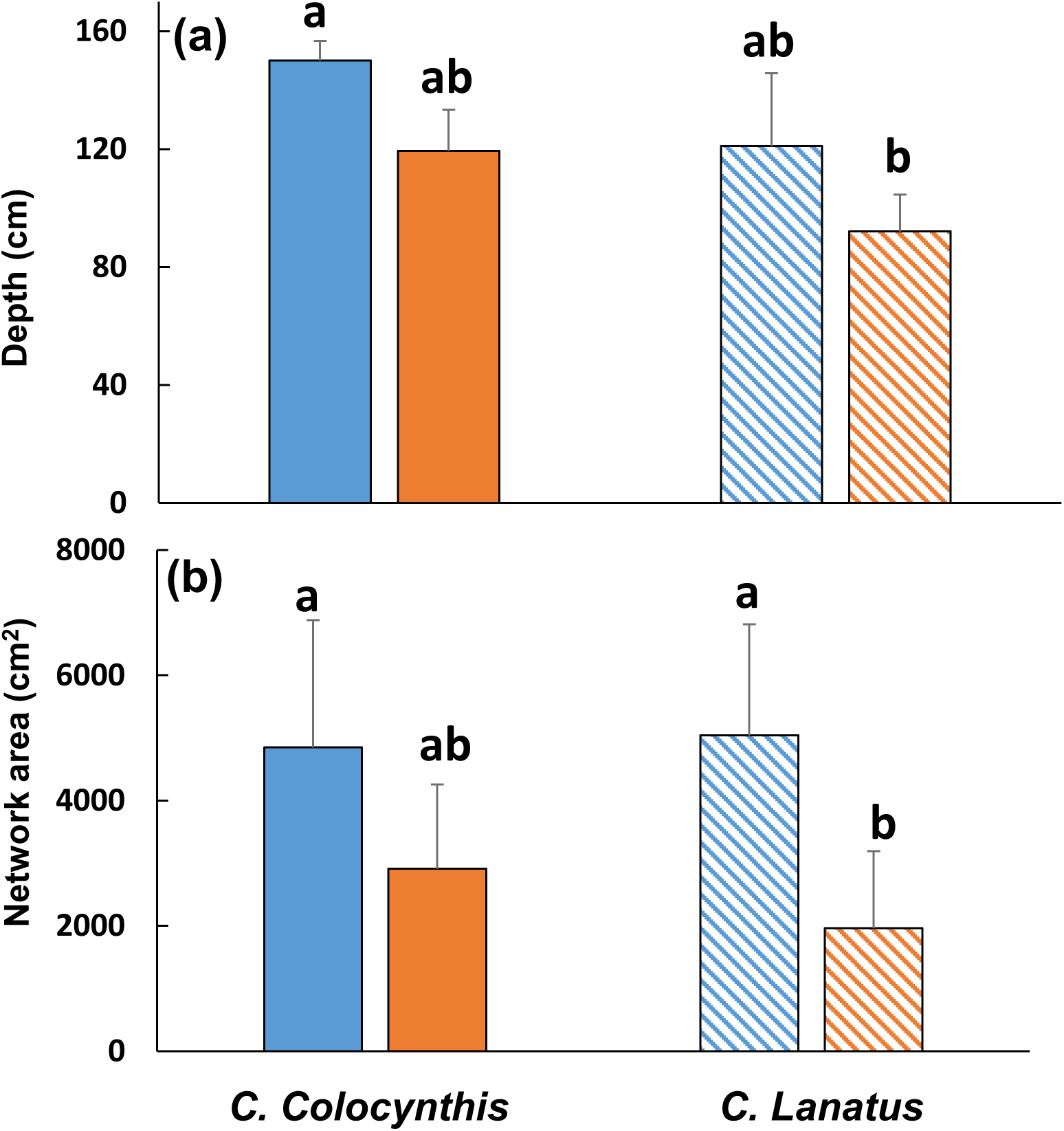
The effect of watermelon genotype and irrigation treatment (Control – 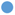, drought – 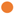) on the roots (a) depth and (b) network area in 10 litter pots. Each genotype*irrigation treatment was repeated four times (beside the *C. lanatus-*Control: *n*=3), bars represent the mean+SD. Different letters showing statistical difference between treatments according to Tukey-HSD test (α<0.05).

## 4. Discussion

Numerous traits associated with drought tolerance have been lost in cultivated crops over the course of evolution and domestication (Roucou et al., 2018; Martín-Robles et al., 2019; Nimmo et al., 2023). Canopy-related adaptations to drought stress have often been deliberately selected against because of their negative associations with yield components (Puli et al., 2024). In contrast, the root system represents a largely unused prospect for enhancing drought tolerance, as many root traits can mitigate water stress without substantially compromising yield (Freschet et al., 2021; Bacher et al., 2022). In this study, we demonstrate that the contrasting drought responses of wild versus domesticated watermelon are primarily driven by differences in their root system architecture.

Root plasticity is a critical function in abiotic stress tolerance, specifically regarding drought. It allows the plant to bypass environmental limitations by adapting its root system architecture and spatial distribution to maintain resource acquisition (Gruber et al., 2013; Karlova et al. 2021; Govta et al., 2025). Different works demonstrated the loss of root system plasticity under drought conditions over the domestication process (Bacher et al., 2023; Wild et al., 2024). This work results demonstrated that while domesticated watermelon root system was inert to water limitation, the wild watermelon root system was adjusted to changing SWC. By increasing its root surface area (SA) in deeper soil layers under drought conditions, the wild watermelon was able to evade water stress. Notably, this expansion of deeper SA did not involve a corresponding increase in biomass allocation, but was achieved specifically through the proliferation of fine roots in the lower soil profile. Recent research suggests that fine roots in deep soil layers are essential for drought resilience across diverse ecosystems (Kwatcho Kengdo et al., 2025).

It is well known that fine roots are the main component in water and nutrient absorption in the soil (Mccormack et al., 2015; Barberon et al., 2016; Cochavi et al., 2020). Their short life span and low carbon cost turn their production to quick and efficient way to explore for water and nutrients in the different soil layers (Mccormack et al., 2015; Cochavi et al., 2018). By producing the largest portion of the root system SA, the fine roots are the key trait to cope with water limitation in both watermelon genotypes. In both species, a significant reduction in root SA was observed and accompanied by a decrease in the average root diameter, although this reduction was more pronounced in the domesticated watermelon. Decreasing root diameter facilitates more efficient water absorption under water-limited conditions (Wasson et al., 2012; Cochavi et al., 2018).

The dynamic relationship between SWC and root SA highlights the fundamental divergence between the irrigation-dependent domesticated watermelon and its wild, desert-origin counterpart. While the domesticated watermelon demonstrated a moderate decrease in root SA as SWC declined, the response of the desert watermelon was significantly more pronounced, with *α* value that is 2.5 times higher. This result underscores that the wild watermelon functions as an ephemeral desert plant, adapted to rapidly exploit transient SWC (Tariq et al., 2024). Conversely, although the domesticated watermelon originated in xeric regions, it has been selected over generations for cultivation under consistent water supplies; consequently, it exhibits a much slower and less plastic response to fluctuations in SWC (Paris, 2015).

Shifting the S\R ratio toward below-ground biomass can assist plants in maintaining physiological function during drought stress (Bloom et al., 1985; Bacher et al., 2022). However, neither watermelon genotype followed this conventional rule. The domesticated watermelon S\R ratio shifted toward the foliage under drought conditions. Conversely, the wild watermelon S\R ratio remained consistently low (close to 1.0) under both control and drought treatments. Rainfall abundance is known to significantly influence S\R ratios (Mokany et al., 2006); thus, the hyper-arid origin of the wild watermelon may explain its inherently high investment in roots. The lack of variation in this ratio under changing soil water content (SWC) indicates a high degree of phenotypic rigidity in the wild genotype.

While many studies emphasize root depth as a primary trait for drought tolerance (Uga et al., 2013; Bacher et al., 2023), our findings suggest a more nuanced reality for watermelon. In the current work, the wild genotype indeed exhibited a deeper root system under control conditions. However, under drought stress, root system depth was reduced in both genotypes. Furthermore, while the total root system dimensions decreased under drought, the reduction in the wild watermelon’s network area was not statistically significant. These results indicate that absolute root depth may not be the primary trait governing drought tolerance in watermelon.

Strict water regulation through stomatal closure is a well-established drought tolerance trait (Blum, 2009; Xu et al., 2016; Guerrieri et al., 2019; Kang et al., 2021). Despite the divergent nature of the two genotypes, both their WUE and iWUE were similar under water-limited conditions. However, the maximal photosynthetic potential (*An*_max_) of the domesticated watermelon under sufficient water supply was significantly higher, highlighting its adaptation as a horticultural crop bred for high productivity. Conversely, while the wild watermelon remained tolerant to low soil water content (SWC), high SWC led to a reduction in Y(II). This suggests that the wild watermelon, given its xeric origin, exhibits a physiological sensitivity to high SWC (e.g. water logging; Zait et al., 2024). The *ETR\An* ratio has been proposed as a diagnostic tool to evaluate the efficiency of electron use for carbon assimilation (Perera-Castro & Flexas, 2023). While the wild desert watermelon maintained a stable *ETR\An* response across all SWC levels, the domesticated variety exhibited higher efficiency only under optimal SWC. The sharp increase in the *ETR\An* ratio of the domesticated variety under water limitation underscores its physiological sensitivity. These divergent responses demonstrate a fundamental shift in watermelon life-history strategies during domestication: transitioning from a conservative strategy adapted to xeric environments to a "risky" strategy optimized for high productivity under sufficient water supply (Guadarrama-Escobar et al., 2024; Lupo & Moshelion 2024). The physiological analysis demonstrated that the wild watermelon adapts its foliage to drought through photosynthetic adjustments rather than through stomatal water regulation (Eppel et al., 2014).

Both genotypes did not change their morphology under drought stress, root and foliage. However, the differences demonstrated the adaption of wild watermelon to drought conditions. Thicker leaf, and specifically cuticle, assist the plant to prevent water loss through the leaf surface (Akram et al., 2023; Aneja et al., 2025). Similarly, the main root area was larger in the desert watermelon. This difference was not an outcome of a larger stele for water transport as by thicker cortex. Previous works point out the importance of large cortex in drought tolerance. Due to their low energetic cost, roots with large cortex area can explore deeper soil layers efficiently, and maintain high physiological functioning of the foliage under drought conditions (Chimungu et al., 2014).

## 5. Conclusions

This study aimed to characterize the drought tolerance traits of the wild watermelon (*C. colocynthis*) and how these were altered through cultivation in the domesticated watermelon (*C. lanatus*). We find that the primary source of drought tolerance in the wild watermelon is efficient root system plasticity (Figure 11). With minimal carbon investment and an adjusted primary root structure, the wild watermelon is able to avoid drought stress through the effective exploitation of deeper soil layers. This trait, however, has been lost in the domesticated watermelon. The lack of root system plasticity in the domesticated watermelon could be due to changes in hormonal signaling or differences in genome structure, but this requires further research.

**Figure 11.**
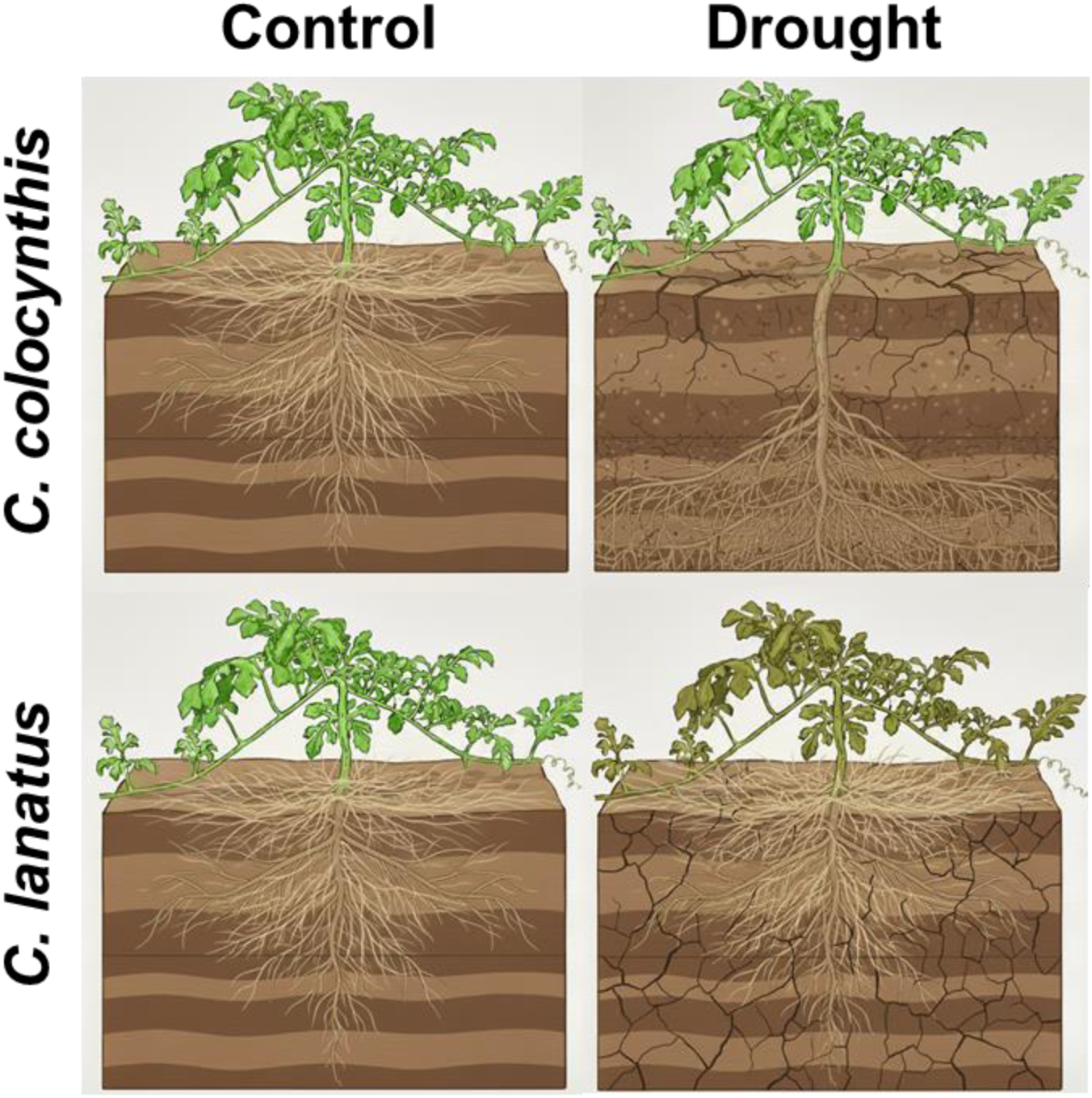
Graphical summary of root system dynamic of *C. lanatus* versus *C. colocynthis* under irrigation and drought conditions. Image were created using Nano Banana (Gemini).

While canopy-related drought traits were often excluded from breeding programs due to their association with reduced yield, root traits associated with drought tolerance may also have been unintentionally lost. However, these traits hold significant potential for improving drought resilience in cultivated crops without compromising yield. Unlike canopy adaptations that primarily reduce water loss through stomatal regulation and leaf area reduction, root-based traits focus on enhancing water uptake efficiency under limited soil moisture conditions as shown in the current work. Future integration of this trait into watermelon cultivars should improve their drought tolerance with minimal yield loss.

## Authors contribution

OES conducted the experiments and collect the data, ZMB conducted the histological analysis, GP conduct the 3-D scanning of the root system, AL assist in conceiving the study, manuscript writing and editing, AC analyzed data and wrote the manuscript.

## Acknowledgment

We wish to thank to our lab technicians Elad Fein and Ofir Tal, for their assistance in the experiments maintenance.

## Data availability statement

The data that support the findings of this study are available from the corresponding author upon reasonable request.

## AI generative statement

During the preparation of this work, the authors employed the Gemini AI agent (Nano-Banana) specifically for the generation of the summary figure (Figure 11). Beyond the creation of this figure, no generative AI or AI-assisted technologies were utilized in the drafting or editing of the manuscript.

