## supplemental figures for "Domestication reduces drought tolerance in watermelon through loss of root plasticity traits"

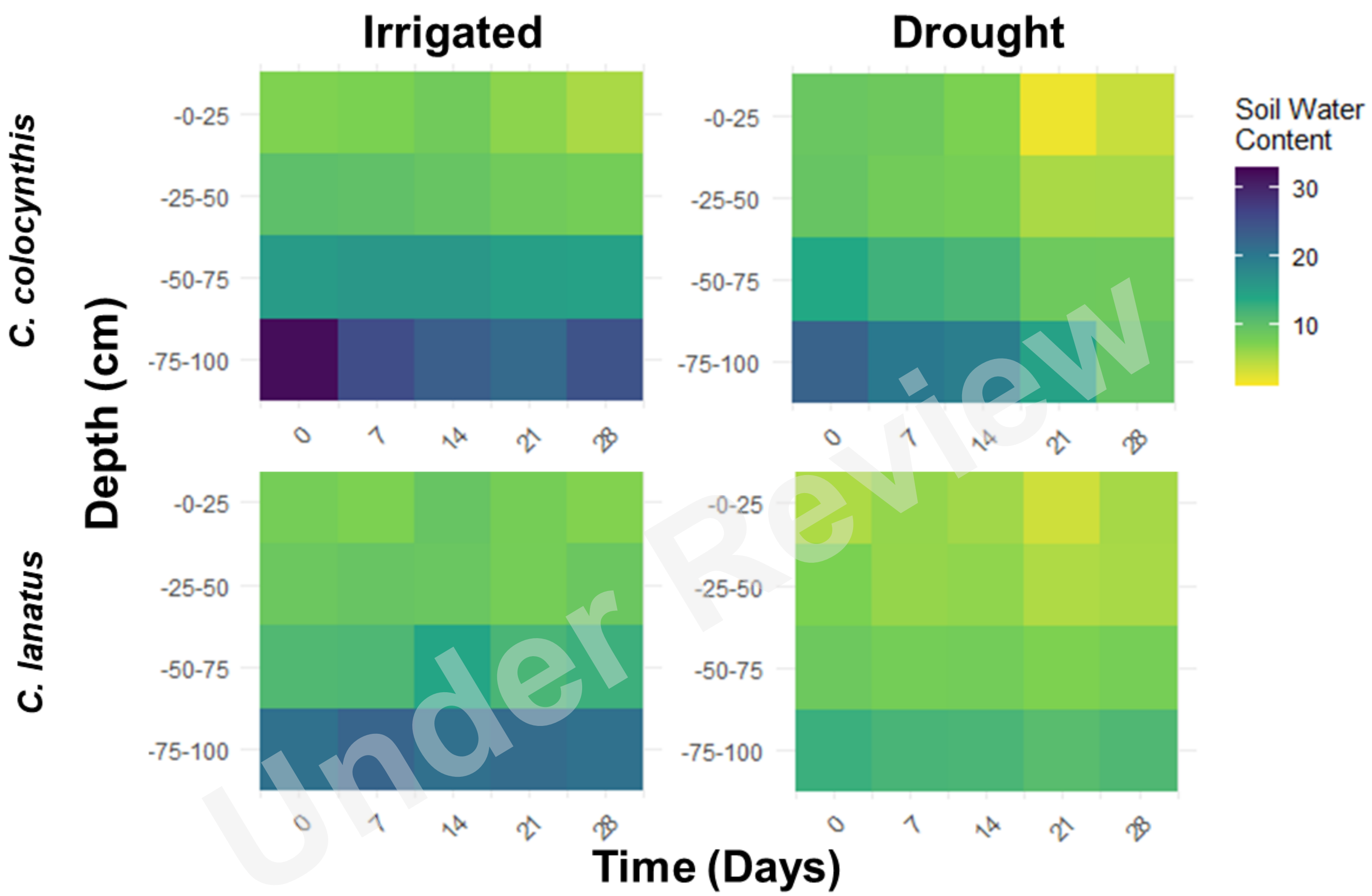


**Figure S1.** Relative water content during the tube experiment in different soil depths (*n*=3).

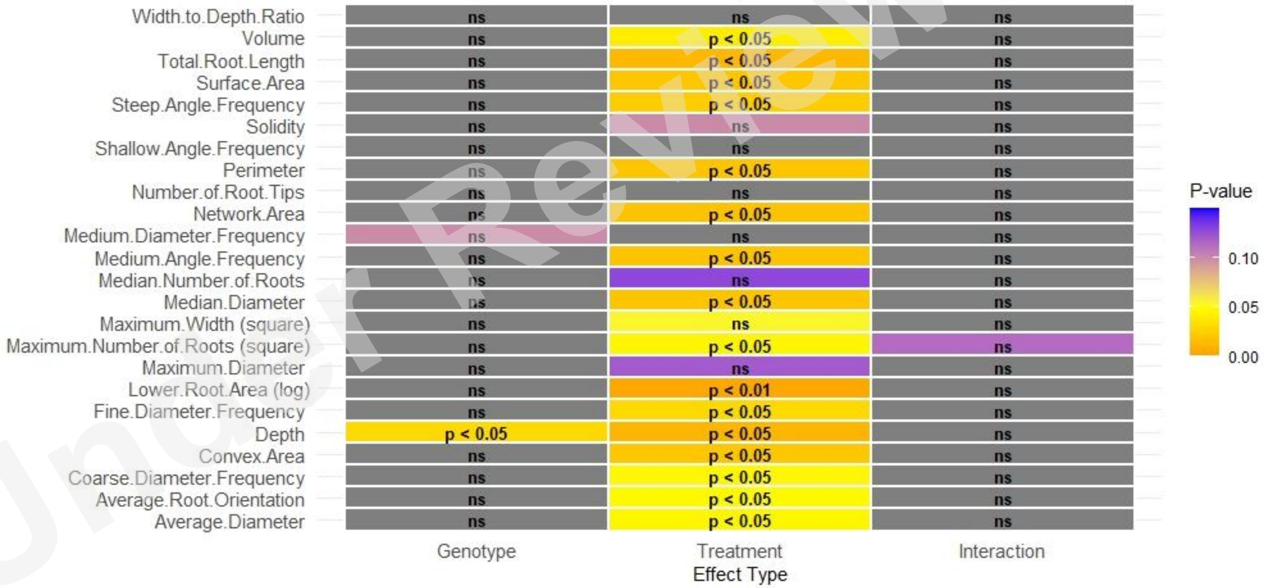


**Figure S2**. Statistical analysis summary for measured variables in the PhenoRoot® system. Parameter with parantces ( ) point on data transformation of the data due to lack of normality or humogenity.

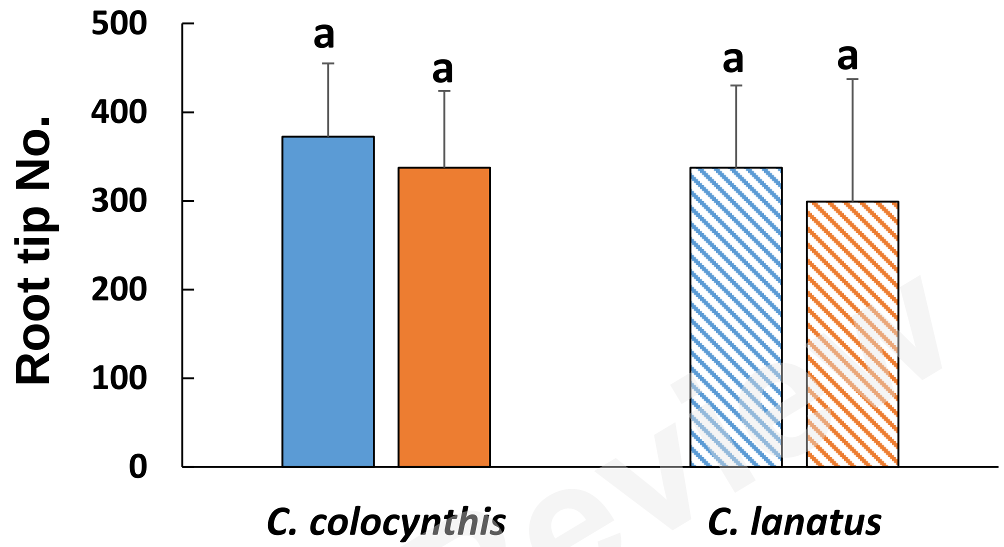


**Figure S3.** Three-dimensional scans and architectural analysis of domesticated (*C. lanatus*) and wild watermelon (*C. colocynthis*) root systems. The effect of irrigation treatment (Control - ●, drought - ●) is shown for root tip number. Root systems were scanned and analyzed using the PhenoRoot® system. Bars represent the mean ± SD.

Under Review
